# Viime-Path: An Interactive Metabolic Pathway Generation Tool for Metabolomics Data Analysis

**DOI:** 10.1101/2023.03.07.531550

**Authors:** Jeff Baumes, Thomas M. O’Connell

## Abstract

**Introduction:** A primary objective of metabolomics studies is identifying the critical metabolic pathways that are altered under specific conditions. There are a number of tools such as enrichment or over-representation algorithms which can identify pathways, but tools for visualizing these pathways beyond static diagrams are limited.

**Objectives:** Development of an open-source software tool to enable users to generate metabolic pathway diagrams that can be interactively adjusted to allow for the most lucid and meaningful representation of the data. The ability to input and integrate multiple metabolic pathways and interactively adjust could facilitate the identification of novel metabolic pathways.

**Methods:** The Viime-Path software package utilizes the metabolic pathways in the Reactome Database. The initial pathway layouts are generated using the Human-like Orthogonal Network Layout (HOLA) algorithm which was designed to take such datasets and map them onto orthogonal network layouts using aesthetic principles derived from the analysis of human-generated networks. The software is written in TypeScript and Vue, utilizing the WebCola graph layout library and D3.js for visualization primitives.

**Results:** We have created an easy-to-use, web-based, open-source tool to easily import metabolic pathways from the Reactome database. The tool allows for interactive optimization of pathways to best represent the most salient features for a given analysis. The tool is available on GitHub at https://github.com/girder/viime-path.

**Conclusions:** The Viime-Path tool will enable the metabolomics community to interactively draw metabolic pathway networks with options to adjust specific components and layout, thus providing new levels of clarity and focus.

## Introduction

A primary objective of metabolomics studies is to take a list of altered metabolites and identify the specific metabolic pathways in which those metabolites participate. There are a number of ways to identify specific pathways, including the use of enrichment or over-representation analyses which can take such lists of metabolites and generate ranked lists of metabolic pathways that are likely to be involved (Booth, Weljie, Turner 2013; Chagoyen, Pazos 2011; Creixell, Reimand, Haider, Wu, Shibata, Vazquez, Mustonen et al. 2015; Gehlenborg, O’Donoghue, Baliga, Goesmann, Hibbs, Kitano, Kohlbacher et al. 2010; Khatri, Sirota, Butte 2012; Misra, van der Hooft 2016; Xia, Wishart 2010); also see useful review by Marco-Ramell et al. (Marco-Ramell, Palau-Rodriguez, Alay, Tulipani, Urpi-Sarda, Sanchez-Pla, Andres-Lacueva 2018). Once the pathways of interest are identified, the investigator is often presented with sets of manually drawn and curated network maps representing these pathways. Such pathways are indeed quite helpful but are limited by fixed graphical representations. The layout and level of detail in these maps were decided by the original teams that drew these pathways, and this representation may not be ideally suited to all studies. **Figure 1** shows the metabolic pathway diagrams for the glycolysis pathway taken from the KEGG (Kanehisa, Sato, Kawashima, Furumichi, Tanabe 2016) and Reactome databases (Creixell, Reimand, Haider, Wu, Shibata, Vazquez, Mustonen et al. 2015; Jassal, Matthews, Viteri, Gong, Lorente, Fabregat, Sidiropoulos et al. 2020). Both of these diagrams are very information-rich, but with that comes a cost in complexity and readability. In the Reactome representation, there are nodes for common pathway features such as protons, water, and carbon dioxide, which may not be of interest in many studies. Other metabolic reaction features such as the transformation of ATP to ADP or reduction of NAD+ to NADH (and the reverse reactions) may be of interest, but often these features are well understood, and they just provide additional nodes that lead to a more crowded pathway. The ability to interactively edit such metabolic features would lead to significantly simplified pathway representations, while maintaining the critical information required for a given analysis.

**Fig 1.**
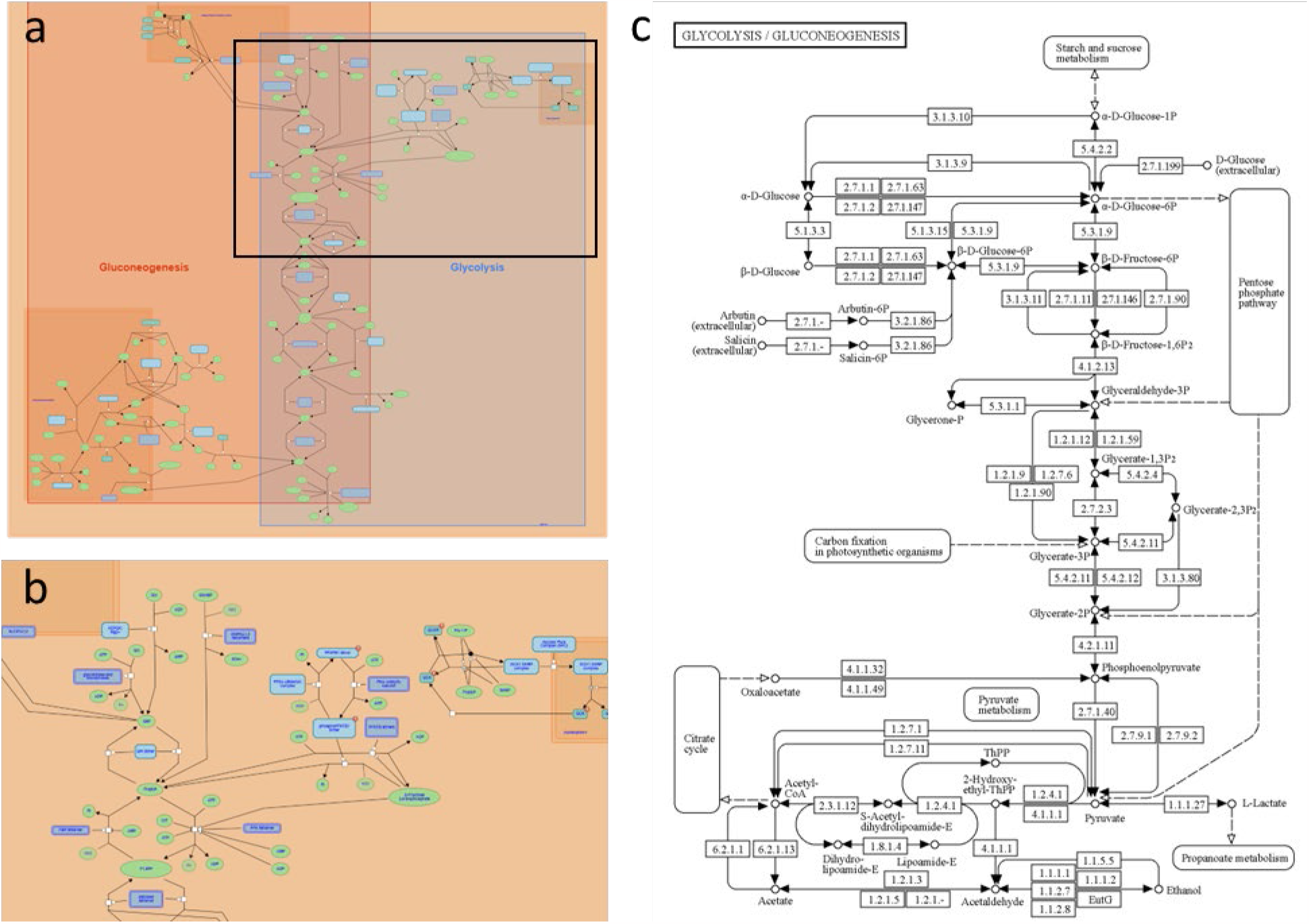
Representations of the glycolysis pathway from the Reactome and KEGG Pathways. **a** Initial Reactome pathway includes both glycolysis and gluconeogenesis. **b** Expansion of a small region of this pathway demonstrates the complexity and level of detail. **c** KEGG pathway representation of glycolysis. Boxes with numbers contain hyperlinks to KEGG database entries for the cellular entities e.g., enzymes that carry out specific reactions.

Manually drawn metabolic pathways inherently take advantage of the guiding principles of graphic design and human visual perception. There is a significant body of literature on the development of algorithms that can draw network diagrams that yield results with the same aesthetic results as those drawn by humans (Dwyer, Lee, Fisher, Quinn, Isenberg, Robertson, North 2009; Purchase, Pilcher, Plimmer 2012; van Ham, Rogowitz 2008). In the context of structured network diagrams such as metabolic pathways, those guiding aesthetic principles were studied in a paper by Kieffer, et al. (Kieffer, Dwyer, Marriott, Wybrow 2016). In this study, a set of volunteers were presented with 8 randomly arranged graphs. Using a custom editing tool, the volunteers could manually adjust the graphs until they “looked good.” Analysis of the results led to the identification of nine specific aesthetic features to guide the network layout, including (1) external placement of tree-like features (2) aesthetic bend points, i.e., nodes do not always have to be placed at grid corners (3) compactness (4) grid-like placement (5) symmetry (6) minimal edge crossings (7) minimal edge bends (8) minimal edge lengths and (9) minimal stress i.e. minimal number of paths through a single node. Using these features as guides, they developed the Human-like Orthogonal Network Layout (HOLA) algorithm. The study found that the graphs drawn using the HOLA algorithm were typically scored as high as the best of the human-drawn graphs. In our pursuit of an interactive software tool to draw metabolic networks, the HOLA algorithm provides an ideal starting point for metabolic networks that can be subsequently optimized by the user.

Viime-Path is designed to provide an interactive pathway mapping tool based on the aesthetic principles of the HOLA algorithm starting with the metabolic pathway information in the Reactome database. The implementation of the HOLA network optimization algorithm was ideally suited to the Reactome database, as it is structured as a graph database with metabolites, proteins, and genes being represented by nodes and edges in a graph (Fabregat, Korninger, Viteri, Sidiropoulos, Marin-Garcia, Ping, Wu et al. 2018). In contrast to the relational database structure, this new format provides more rapid and efficient queries. With Viime-Pathways, users can query different metabolic pathways and automatically generate a metabolic network map. Once an initial map is generated, the user may delete specific features as desired and interactively move the nodes to generate new maps that maintain the guiding aesthetic principles of the HOLA algorithm. Visualization options also include adjustment of node size and shape, size of font for labels, and background coloring to show cellular compartments and highlight specific metabolites of interest. Additionally, this tool will allow the import of multiple metabolic pathways so that more complex, interconnected metabolic pathways can be examined.

Currently Viime-Path is a standalone tool, but the ultimate goal is for this to be integrated as a module in the Viime program; www.viime.org; (Choudhury, Beezley, Davis, Tomeck, Gratzl, Golzarri-Arroyo, Wan et al. 2020)). Viime is an open-source, web-accessible program that was developed to facilitate the integration, analysis, and visualization of metabolomics data from multiple analytical platforms.

## 1. Materials and Methods

### 1.1 Software and access

Viime Path makes heavy use of Reactome data to query and construct its networks. The Reactome database is a curated collection of pathways and their associated reactions, proteins, and metabolites for humans and other species. Reactome provides programmatic access to querying and retrieving pathways and related domain objects via their Content Service REST API documented at https://reactome.org/ContentService/. This API enables Viime Path to be implemented as a pure browser-based application without the need for a dedicated server.

The code is structured as a Vue component composed of a set of reactive controls on the left panel and a main visualization area. Styling for the application utilizes the CSS libraries Tailwind and DaisyUI. Metabolites of interest and pathway search controls dynamically fetch relevant information from Reactome based on user queries. Based on the user’s selection of pathways, the application generates an HOLA orthogonal graph layout using the WebCola library and renders it with D3 with a full pan and zoom interface. The user may selectively show, hide, or highlight individual pathways, metabolites, and reactions using the side panel and by interacting with the graph itself, which utilizes Vue’s reactivity to update the view. Network nodes may also be manually positioned. Nodes may be colored by compartment (e.g., cytosol vs. mitochondria) or by pathway. In addition, the background may reflect either of these two facets by coloring the cells of a Voronoi diagram with points located at network nodes.

The code is available at https://github.com/girder/viime-path and a live instance of the application is deployed to https://girder.github.io/viime-path/.

### 2.2. Operation of Pathway Editor

The process of generating an interactive pathway diagram starts with loading a specific metabolic pathway or pathways into the tool. This is done by directly inputting a Reactome Pathway identifier number or via a keyword search. A hyperlink to the Reactome Pathway Browser can be used to search for pathways via the Reactome interface. Pathways in Reactome are organized in a hierarchical manner such that major pathway descriptions such as “Metabolism of Carbohydrates” have numerous sub-pathways below them. In this case, those would include pathways such as Glucose metabolism, Glycogen metabolism, Fructose metabolism and more. Each pathway description has a unique identifier (ID) which can be loaded into the Viime-Path Pathway search box. If the selected pathway has sub-pathways all of them will be loaded. This may result in an excess of information and a highly complicated layout. Users may want to drill down to lower pathway levels in the interest of simplicity. When using the keyword search, some queries will return several pathways and the user can select one or all of them. For example, “glycolysis” will yield two pathways (1) Glycolysis and (2) Regulation of glycolysis by fructose.

A list of metabolites can be entered into the Metabolites of Interest box so that they are highlighted on the map with a red node border. The list can be composed of common names or identifiers such as HMDB or ChEBI numbers. For some metabolites, inputting a single name or ID can yield multiple metabolites. This comes about for two reasons. First, the search may yield multiple metabolites that contain that same root word. For example, searching for glucose will also yield glucose-6-phosphate along with other metabolites that include glucose in the name. Second, the Reactome database has metabolites listed separately by compartment such that one metabolite will have separate entries for its location in each compartment. For example, adding the metabolite lactate yields separate results for cytosolic and mitochondrial lactate. The specific metabolites to be highlighted can be selected from the retrieved metabolite list.

The appearance of the map can be modified in a number of different ways. The initial layout of the metabolic network was drawn using the HOLA algorithm as the starting point. The metabolite nodes are colored gray by default and can be colored by compartment. The reaction nodes are white by default and can be colored by pathway. The background color can be a uniform white or different regions can be colored according to the compartment or pathways of the metabolites and reactions. The node size and shape can be adjusted with the shape transitioning from a square to a circle. The font size for the node labels can also be adjusted. Lastly, the spacing of the map can be adjusted to make the map more compact or expanded.

A key feature of Viime-Path is the ability to remove specific nodes to create less crowded, simplified pathway maps. Left clicking on a metabolite node gives options to (1) hide the node (2) split the node and (3) hide the label. If a node is hidden, the node is no longer visible, and the map is redrawn to adjust to the newly simplified network. Note that all instances of that particular node are removed. For example, if NH4+ is selected then all of the NH4+ nodes in that compartment will be removed and the map redrawn. Another way to make a map less crowded is to split a node. When a metabolite has multiple connections, this metabolite can be split so that each connection goes to a separate node. When nodes are hidden or split, they are listed on the left panel and can be selectively re-integrated into the map if desired.

Nodes can be dragged from one location to a new one with the new location optimized according to the HOLA algorithm. Each movement causes an optimization of the entire map, but adjustments tend to be local, i.e., not leading to rearrangements of the network distal to the deleted node. By default, the metabolite nodes are labeled, and the reactions are not. This is because the reaction names can be quite long, leading to a cluttered map. By clicking on any node, metabolite or reaction, the label can be turned on or off to highlight specific features. After all of the interactive adjustments to the map have been made, it can be exported as a png or svg file for use in publications, grant proposals, etc.

## 2. Results

Viime-Path was used to automatically generate the initial pathway diagram for glycolysis shown in **Figure 2A**. This diagram contains all of the nodes that are included in the Reactome database entry for Glycolysis, including nodes representing reaction elements such as H+, CO2, ATP, AMP, and NAD(H). **Figure 2B** shows the diagram after many of these nodes have been removed. This much simpler diagram can be further optimized by moving around the nodes to generate a pathway map that best represents the important features of the pathway. The expansion shown in **Figure 2C** shows the grid-like, simplified representation of the one part of the cytosolic component of the glycolysis pathway.

**Fig. 2.**
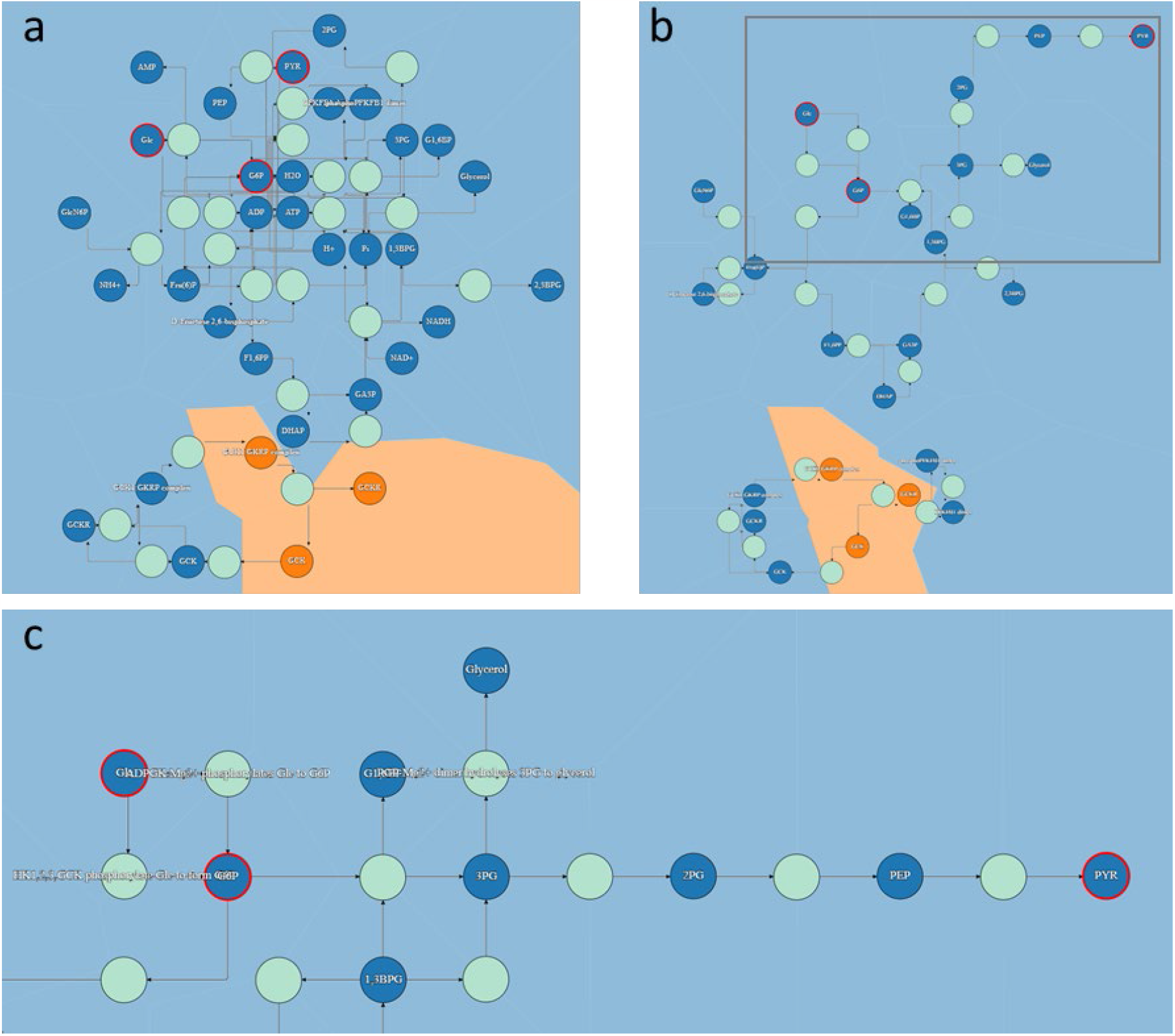
Viime-Path generated pathway maps for glycolysis. **a** Initial map of the glycolysis pathway with the metabolites glucose, glucose-6-phosphate and pyruvate highlighted with a red node border. **b** Simplified map after hiding cytosolic NAD+, NADH, H+, ATP, ADP, Pi, H2O and NH4+. **c** Expansion of a specific region of the pathway indicated by the gray box in b, after moving nodes to clarify the one of the pathways from glucose to pyruvate. Labels for several enzymes were turned on including ADP-specific glucokinase, Hexokinase 1 and Phosphoglycolate phosphatase (also known as glycerol-3-phosphate phosphatase).

## 3. Conclusions

Viime-path provides a novel way to interactively generate customized metabolic pathways using an algorithm designed to mimic the aesthetic principles that guide human-drawn pathways. By enabling the researcher to interact with the data in this way, we hope that it will facilitate new insights into metabolism that would not arise by looking at a static pathway map. The ability to upload and manipulate multiple metabolic pathways at a time could lead to the identification of novel connections between pathways and potentially novel pathways. Beyond research, we anticipate that Viime-Path could be a valuable teaching tool whereby students can generate pathway maps and interactively manipulate them to learn the flow of metabolites from one end of the pathway to the other.

## Author Conflicts of Interest

Jeff Baumes and Thomas M. O’Connell declare no conflicts of interest.

## Author Contributions

Jeff Baumes developed and implemented all of the software tools and performed all of the coding. Thomas M. O’Connell conceived of the overall design of the software and wrote the manuscript. Both authors read and approved the manuscript.

## Compliance with Ethical Standards

This article does not contain any of the studies with human and/or animal participants performed by the author.

## Software Availability

The Viime-Path software tool is available at https://github.com/girder/viime-path

## Statements and Declarations

JB is an employee of Kitware Inc., a scientific visualization software company. TMO has no competing interests.

## Notes

https://girder.github.io/viime-path/

### Competing Interest Statement

The authors have declared no competing interest.

## References

Booth, S. C., A. M. Weljie, R. J. Turner (2013). Computational tools for the secondary analysis of metabolomics experiments. Comput Struct Biotechnol J 4, e201301003 doi:10.5936/csbj.201301003

Chagoyen, M., F. Pazos (2011). MBRole: enrichment analysis of metabolomic data. Bioinformatics 27, 730–1 doi:10.1093/bioinformatics/btr001

Choudhury, R., et al. (2020). Viime: Visualization and Integration of Metabolomics Experiments. J Open Source Softw 5, doi:10.21105/joss.02410

Creixell, P., et al. (2015). Pathway and network analysis of cancer genomes. Nat Methods 12, 615–621 doi:10.1038/nmeth.3440

Dwyer, T., et al. (2009). A comparison of user-generated and automatic graph layouts. IEEE Trans Vis Comput Graph 15, 961–8 doi:10.1109/TVCG.2009.109

Fabregat, A., et al. (2018). Reactome graph database: Efficient access to complex pathway data. PLoS Comput Biol 14, e1005968 doi:10.1371/journal.pcbi.1005968

Gehlenborg, N., et al. (2010). Visualization of omics data for systems biology. Nat Methods 7, S56–68 doi:10.1038/nmeth.1436

Jassal, B., et al. (2020). The reactome pathway knowledgebase. Nucleic Acids Res 48, D498–D503 doi:10.1093/nar/gkz1031

Kanehisa, M., Y. Sato, M. Kawashima, M. Furumichi, M. Tanabe (2016). KEGG as a reference resource for gene and protein annotation. Nucleic Acids Res 44, D457–62 doi:10.1093/nar/gkv1070

Khatri, P., M. Sirota, A. J. Butte (2012). Ten years of pathway analysis: current approaches and outstanding challenges. PLoS Comput Biol 8, e1002375 doi:10.1371/journal.pcbi.1002375

Kieffer, S., T. Dwyer, K. Marriott, M. Wybrow (2016). HOLA: Human-like Orthogonal Network Layout. IEEE Trans Vis Comput Graph 22, 349–358

Marco-Ramell, A., et al. (2018). Evaluation and comparison of bioinformatic tools for the enrichment analysis of metabolomics data. BMC Bioinformatics 19, 1 doi:10.1186/s12859-017-2006-0

Misra, B. B., J. J. van der Hooft (2016). Updates in metabolomics tools and resources: 2014-2015. Electrophoresis 37, 86–110 doi:10.1002/elps.201500417

Purchase, H. C., C. Pilcher, B. Plimmer (2012). Graph Drawing Aesthetics-Created by Users, Not Algorithms. IEEE Trans Vis Comput Graph 18, 81–92 doi:10.1109/TVCG.2010.269

van Ham, F., B. E. Rogowitz (2008). Perceptual organization in user-generated graph layouts. IEEE Trans Vis Comput Graph 14, 1333–9 doi:10.1109/TVCG.2008.155

Xia, J., D. S. Wishart (2010). MSEA: a web-based tool to identify biologically meaningful patterns in quantitative metabolomic data. Nucleic Acids Res 38, W71–7 doi:10.1093/nar/gkq329

